# The Stochastic Resonance model of auditory perception: A unified explanation of tinnitus development, Zwicker tone illusion, and residual inhibition

**DOI:** 10.1101/2020.03.27.011163

**Authors:** Achim Schilling, Konstantin Tziridis, Holger Schulze, Patrick Krauss

**Affiliations:** Neuroscience Lab, Experimental Otolaryngology, Friedrich-Alexander University Erlangen-Nürnberg (FAU), Germany; Cognitive Computational Neuroscience Group at the Chair of English Philology and Linguistics, Friedrich-Alexander University Erlangen-Nürnberg (FAU), Germany; FAU Linguistics Lab, Friedrich-Alexander University Erlangen-Nürnberg (FAU), Germany; Department of Otorhinolaryngology/Head and Neck Surgery, University of Groningen, University Medical Center Groningen, The Netherlands

**Keywords:** Auditory Phantom Perception, Somatosensory Projections, Dorsal Cochlear Nucleus, Speech Perception

## Abstract

Stochastic Resonance (SR) has been proposed to play a major role in auditory perception, and to maintain optimal information transmission from the cochlea to the auditory system. By this, the auditory system could adapt to changes of the auditory input at second or even sub-second timescales. In case of reduced auditory input, somatosensory projections to the dorsal cochlear nucleus would be disinhibited in order to improve hearing thresholds by means of SR. As a side effect, the increased somatosensory input corresponding to the observed tinnitus-associated neuronal hyperactivity is then perceived as tinnitus. In addition, the model can also explain transient phantom tone perceptions occurring after ear plugging, or the Zwicker tone illusion. Vice versa, the model predicts that via stimulation with acoustic noise, SR would not be needed to optimize information transmission, and hence somatosensory noise would be tuned down, resulting in a transient vanishing of tinnitus, an effect referred to as residual inhibition.

## Stochastic Resonance

In engineering, the term *noise*, defined as undesirable disturbances or fluctuations, is considered to be the “fundamental enemy” (McDonnell & Abbott 2009) for error-free information transmission, processing, and communication. However, a vast and even increasing number of studies show the various benefits of noise in the context of signal detection and processing. Here, the most important phenomena are called stochastic resonance (McDonnell & Abbott 2009), coherence resonance (Pikovsky & Kurths 1997), and recurrence resonance (Krauss et al., 2019a).

The term stochastic resonance (SR), which has been introduced by Benzi in 1981 (Benzi et al., 1981), refers to the phenomenon that signals otherwise sub-threshold for a given sensor can be detected by adding a random signal, i.e. noise, of appropriate intensity to the sensor input (Gammaitoni et al., 1998; Moss et al., 2004). Figure 1 illustrates this principle.

**Figure 1:**
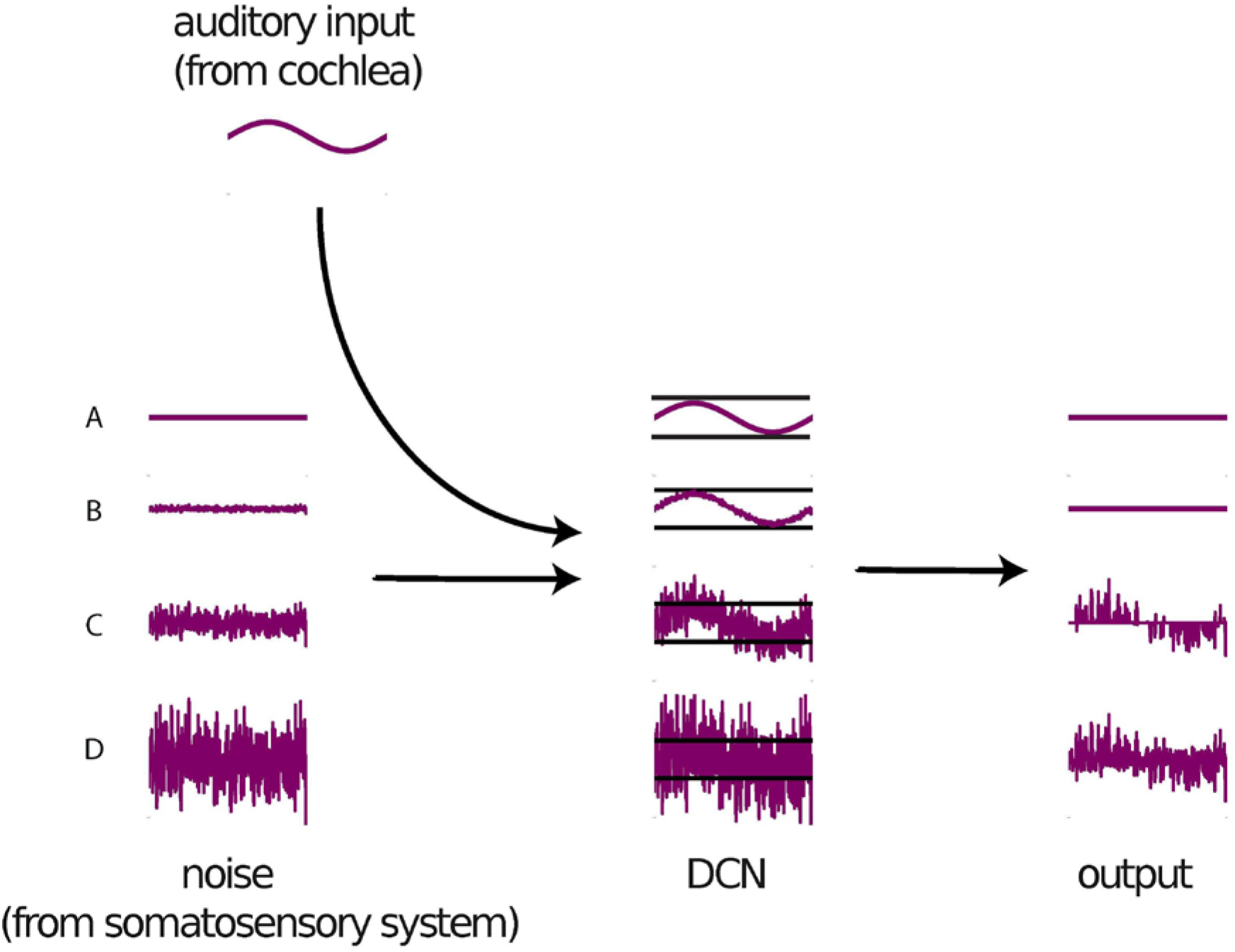
Principle of Stochastic Resonance. The auditory input without any added noise is too weak to pass the threshold (A). Also if the intensity of added noise is too weak, the sum of auditory input and noise cannot pass the threshold (B). Both cases result in zero output. In contrast, if the optimal amount of noise is added to the signal before thresholding, the resulting output’s envelope resembles the auditory input signal (C). However, if the noise intensity is further increased, the signal vanishes again in the noisy output (D).

SR has been found ubiquitously in nature in a broad range of systems from physical to biological contexts (Wiesenfeld & Moss 1995; Hänggi 2002). In particular in neuroscience, SR has been demonstrated to play an essential role in virtually all kinds of systems (Faisal et al., 2008): from tactile (Douglass et al., 1993; Collins et al., 1996), auditory (Mino 2014) and visual (Aihara et al., 2008) perception (Ward et al., 2002), through memory retrieval (Usher & Feingold 2000) and cognition (Chandrasekharan et al. 2005), to behavioral control (Ward et al., 2002; Kitajo et al., 2003). SR explains how the brain processes information in noisy environments at each level of scale from single synapses (Stacey & Durand 2001), through individual neurons (Nozaki et al., 1999; Kosko & Mitaim 2003), to complete networks (Gluckman et al., 1996).

In self-adaptive signal detection systems exploiting SR, the optimum intensity of the noise is continuously adjusted so that information transmission is maximized, even if the characteristics and statistics of the input signal change (Figure 2). For this processing principle, the term adaptive SR has been coined (Mitaim & Kosko 1998, 2004; Wenning & Obermayer 2003; Krauss et al., 2017).

**Figure 2:**
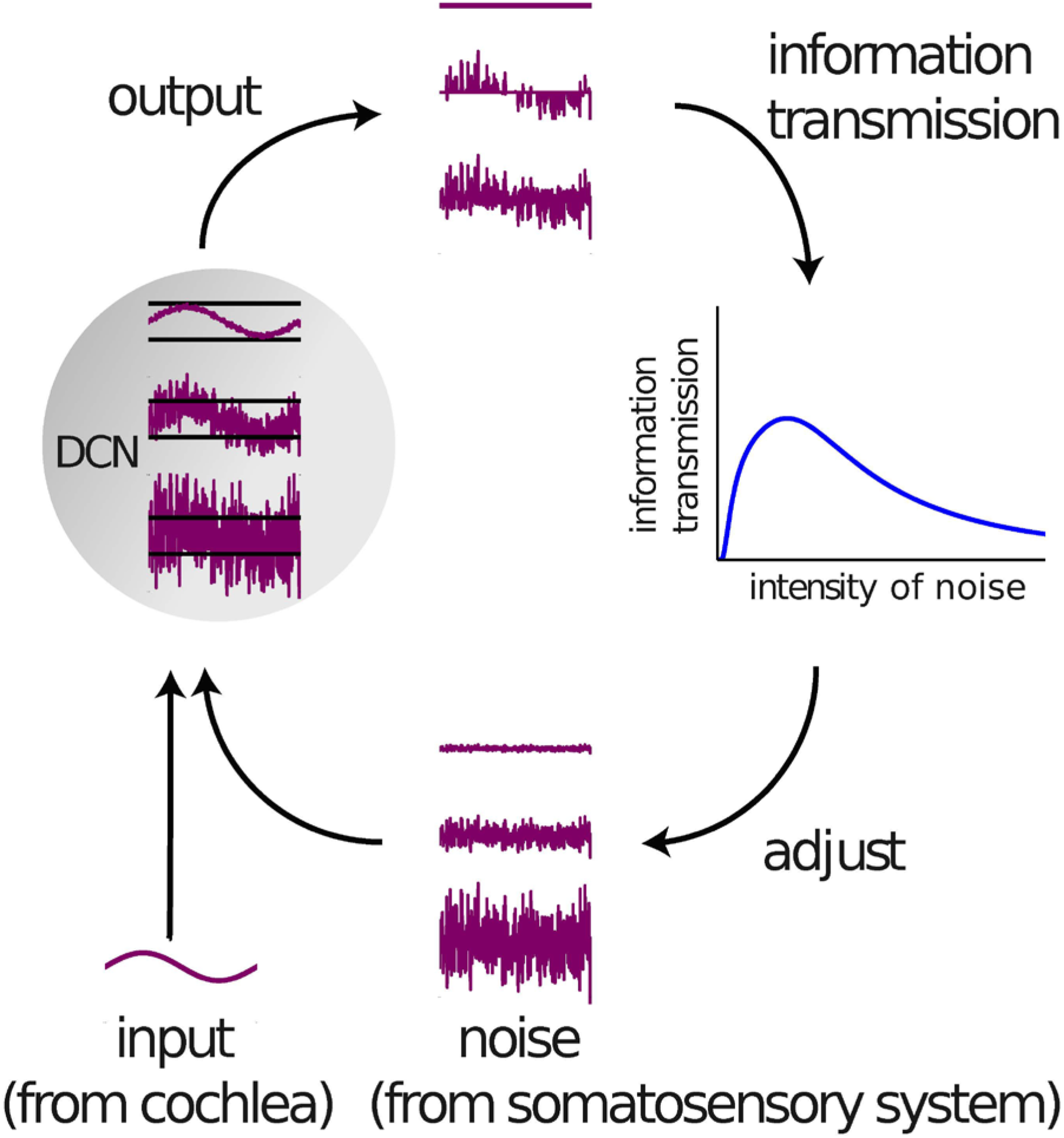
Adaptive Stochastic Resonance control circuit in the DCN. In self-adaptive signal detection systems based on SR, the optimum noise level is continuously adjusted via a feedback loop, so that the system’s response in terms of information throughput remains optimal, even if the properties of the input signal change. In the SR model of tinnitus development, this process takes place in the DCN. The input signal comes from the cochlea, the noise from the somatosensory system.

## Tinnitus Development

In a number of recently published studies, we demonstrated theoretically and empirically that SR might be a major processing principle of the auditory system that serves to partially compensate for acute or chronic hearing loss (Krauss et al., 2016, 2017, 2018, 2019b; Gollnast et al., 2017). According to our model, the noise required for SR is generated within the brain and then perceived as a phantom sound. We have proposed that it corresponds to increased spontaneous neuronal firing rates in early processing stages of the auditory brain stem - a phenomenon which is frequently observed in both humans with subjective tinnitus (Wang et al., 1997; Ahlf et al., 2012; Tziridis et al., 2015; Wu et al., 2016) and animal models, where the presence of tinnitus is tested using behavioral paradigms (Gerum et al., 2019, Schilling et al., 2017, Turner et al., 2006). Furthermore, tinnitus is assumed to be virtually always caused by some kind of either apparent (Heller 2003; König et al., 2006; Nelson & Chen 2004; Shore et al., 2016) or hidden hearing loss (Schaette & McAlpine 2011; Liberman & Liberman 2015). From this point of view, auditory phantom perceptions like tinnitus (or even the Zwicker tone, cf. below) seem to be a side effect of an adaptive mechanism within the auditory system whose primary purpose is to compensate for reduced input through continuous optimization of information transmission (Krauss et al., 2016, 2017, 2018, 2019b). This new interpretation may also explain why auditory sensitivity is increased in tinnitus ears (Hebert et al., 2013; Gollnast et al., 2017): the increased amount of neural noise during tinnitus improves auditory sensitivity by means of SR.

The dorsal cochlear nucleus (DCN) has been shown to be the earliest processing stage where acoustic trauma, including complete cochlea ablation (Zacharek et al., 2002), causes increased spontaneous firing rates (Kaltenbach et al., 1998; Kaltenbach & Afman 2000; Zacharek et al., 2002; Wu et al., 2016). Interestingly, this increase in spontaneous activity, i.e. neural hyperactivity, is correlated with the strength of the behavioral signs of tinnitus in animal models (Kaltenbach et al., 2004). Furthermore, the hyperactivity is localized in those regions of the DCN that are innervated by the damaged parts of the cochlea (Kaltenbach et al., 2002). Gao and colleagues (Gao et al., 2016) recently described changes in DCN fusiform cell spontaneous activity after noise exposure that perfectly supports the proposed SR mechanism. In particular, the time course of spontaneous rate changes shows an almost complete loss of spontaneous activity immediately after loud sound exposure (as no SR is needed due to stimulation that is well above threshold), followed by an overcompensation of spontaneous rates to levels well above pre-exposition rates since SR is now used to compensate for acute hearing loss (Gao et al., 2016).

It is well known that the DCN receives not only auditory input from the cochlea, but also input from the somatosensory system (Ryugo et al., 2003; Shore & Zhou 2006; Wu et al., 2015), and that noise trauma alters long-term somatosensory-auditory processing in the DCN (Dehmel et al., 2008, 2012; Shore 2011, Wu et al., 2016), i.e. somatosensory projections are up-regulated after hearing loss (Zeng et al., 2012). In addition, DCN responses to somatosensory stimulation are enhanced after noise-induced hearing loss (Shore et al., 2008; Shore 2011; Wu et al., 2016). Therefore, we previously proposed the possibility that the neural noise which is necessary for SR is injected into the auditory system via somatosensory projections to the DCN (Krauss et al., 2016, 2018, 2019b), and that these non-auditory projections into the DCN are the cause of the altered “spontaneous activity” within the DCN after hearing loss described previously (Gao et al., 2016). From an information processing point of view, somatosensory inputs are completely uncorrelated, i.e. have no mutual auditory information. Hence, these somatosensory inputs are perfectly suited to serve as a random signal, i.e. noise, in the context of SR, and this seems to be the reason why the auditory system does not generate the noise needed for SR itself.

Our idea that cross-modal SR, with cochlear inputs being the signal and somatosensory projections being the noise (Figure 2), is a key processing principle of the auditory system and actually takes place in the DCN (Krauss et al., 2018) is supported by a large number of different findings. For instance, it is well known, that jaw movements lead to a modulation of subjective tinnitus loudness (Pinchoff et al., 1998). This may easily be explained within our framework, as jaw movements alter somatosensory input to the DCN: Since this somatosensory input corresponds to the noise for SR, auditory input to the DCN is modulated through this mechanism, and the altered noise level would then be perceived as modulated tinnitus (Krauss et al., 2016, 2018, 2019b). Along the same line, one may explain why both, the temporo-mandibular joint syndrome and whiplash, frequently cause so called somatic tinnitus (Levine 1999; Shore et al., 2007).

Furthermore, the finding of Tang and Trussell that somatosensory input and hence tinnitus sensation may also be modified by serotonergic regulation of excitability of principal cells in the DCN (Tang & Trussell 2015, 2017) supports the SR model. It even provides a mechanistic explanation of salicylate induced tinnitus, since salicylate affects DCN processing by disinhibition of somatosensory inputs (Koerber et al.,1966, Stoltzberg et al., 2012). Thus, it increases the noise in the auditory system, which then may again be perceived as a phantom sound.

Finally, and maybe most remarkable, electro-tactile stimulation of finger tips, i.e. increased somatosensory input, significantly improves both, melody recognition (Huang et al., 2020) and speech recognition (Huang et al., 2017) in patients with cochlear implants. Very recently, we were able to reproduce and mechanistically explain this finding, using a hybrid-computational model that exploits SR. The model consists of a cochlea model, a DCN model and an artificial deep neural network trained on a speech recognition task representing all further processing stages of the auditory pathway beyond the DCN. Simulated hearing loss, i.e. weakening the input from the cochlea model to the DCN model, reduced accuracy for speech recognition in the deep neural network, as expected. However, subsequent noise, i.e. somatosensory, input to the DCN model results in an improved accuracy for speech recognition (Schilling et al., 2020).

## Zwicker Tone Illusion

The Zwicker tone effect was discovered by Eberhard Zwicker in 1964 and is a temporal auditory phantom percept which was originally induced by the presentation of a 60 dB broadband noise with a spectral gap (notched noise) with a gap-width of half an octave (Zwicker 1964). The Zwicker tone was described as “Negative Auditory After Image”, although the underlying mechanisms generating an “After Image” are supposed to be different in the visual system. The Zwicker tone perception is not exclusively induced by a notched noise stimulus, but can also be caused by low-pass noise or white noise with a loud pure tone embedded (Fastl et al., 2001, Franosch et al., 2003).

Several models exist trying to explain the Zwicker tone percept. For example, Franosch and colleagues viewed the Zwicker tone as an asymmetric lateral inhibition effect along the auditory pathway (Franosch et al. 2003). In this view, the neurons in the DCN are disinhibited by surrounding neurons, which receive less stimulus driven activity due to the notch.

Another model suggested the Zwicker tone to be caused by a prediction error within the cortex in combination with an increased spontaneous rate of auditory pathway neurons at frequency ranges deprived by the notch within the presented broadband noise (Hullfish et al., 2019). However, these models have certain shortcomings such as they do not account for all properties of Zwicker tone percepts (described in the following) or do not describe the effect on a neuronal network level.

It has previously been proposed that the Zwicker tone and tinnitus and thus also the neural mechanisms of these two auditory phantom perceptions are closely connected (Lummis and Guttmann, 1972, Hoke & Hoke, 1996; Mohan et al., 2020,), and a number of findings support this assumption: For example, Parra and Pearlmutter were able to show that people with a tinnitus percept are also more likely to perceive a Zwicker tone percept (Parra & Pearlmutter, 2007). Additionally, Wiegrebe and coworkers showed that the presence of a Zwicker tone leads to decreased auditory thresholds of 13 dB even in normal hearing subjects (Wiegrebe et al., Norena et al. 1999), a finding which may easily be explained within our above described model of SR, since a similar effect can be observed in tinnitus patients (Gollnast et al., 2017, Krauss et al. 2016) who have improved hearing thresholds in comparison to patients without tinnitus, at least within frequency ranges below 3 kHz. In this context, psychoacoustic experiments revealed that notched noise presentation leads to higher sensitivity to tones embedded in noise (Zhou et al., 2010).

Next, human studies using MEG showed that Zwicker tone perception correlates with a reduced alpha activity (Leske et al., 2014) in the auditory cortex. Interestingly, the effect of reduced alpha activity is also correlated to tinnitus perception (Weisz et al. 2007, Weisz et al. 2011).

Furthermore, in most models tinnitus is supposed to be caused by hearing loss (Moffat et al., 2009) through e.g. cochlea damage or hidden hearing loss which cannot be detected by pure-tone audiograms but is characterized by a deafferentation of the inner hair cells (Liberman & Liberman, 2015; Paul et al., 2017). Analogously, the induction of the Zwicker tone through notched noise can be viewed as a deprivation of certain inner hair cells, that is, a temporary and reversible hearing loss (Hullfish et al., 2019).

These observations and resemblances support the view that the neural mechanisms of Zwicker tone and acute tinnitus are similar and that therefore the Zwicker tone may be a good model for tinnitus (Hullfish et al., 2019, Franosch et. al. 2003, Krauss et al. 2018, Wrzosek et al., 2017, Norena et al., 2000, Norena et al., 2002, Norena et al. 1999). As a result, the investigation of the Zwicker tone has recently attracted further attention. Norena & Eggermont showed that Zwicker tone related neuronal activity changes can be observed on time scales in the range of seconds (Norena & Eggermont, 2003). In particular, cats were implanted with multi-electrode arrays and notched noise stimuli of 1 s duration were presented. It could be shown that neurons in the auditory cortex representing frequencies within the range of the notch show increased firing rates after notched noise presentation (Norena et al., 2003). This result indicates that the Zwicker tone is correlated with a hyperactivity of neurons along the complete auditory pathway that represent the frequency notch, although to our knowledge systematic studies of activity along the auditory pathway in animals during Zwicker tone induction are missing.

Despite all these similarities between the Zwicker tone and acute tinnitus, there are only few mechanistic explanation approaches on a neural network level (Okamoto et al., 2005). Our stochastic resonance model (Krauss et al., 2016, see above) provides such a mechanistic explanation of Zwicker tone percepts. As stated above the presentation of a notched noise stimulus can be viewed as temporary hearing loss or deprivation of inner hair cells located within the frequency notch within the tonotopic gradient (Hullfish et al., 2019; Krauss et al., 2018). According to our model, this reduced input would cause SR within the auditory system to restore hearing by optimizing information transmission at the level of the DCN via increased neuronal noise (as described above). This increase of the neural noise would take place within the frequency channels of the spectral notch, leading to a hyperactivity of the respective neurons in the DCN (Krauss et al., 2016). This hyperactivity is transmitted along the auditory pathway and causes a Zwicker tone percept at the cortical level.

Our explanation is supported by the observation that notched noise stimulation leads to hyperactivity of auditory cortex neurons representing the notch frequency (cf. Norena et al., 2003) via disinhibition (cf. Weisz et al., 2007, Weisz et al. 2011). Furthermore, only the SR mechanism may explain improved hearing thresholds for frequencies near the Zwicker tone frequency during Zwicker tone perception (cf. Wiegrebe et al., 1996, Norena et al. 1999): internal noise from the somatosensory system is increased in the deprived frequency ranges (notch frequency range) in order to compensate for reduced auditory input by means of SR. This, in turn, leads as a side effect to improved hearing thresholds for neighboring frequencies above and below the notch. Additionally, the SR feedback control circuit (Figure 2) operates on time scales in the range of or below a second and thus fits to the observation of Zwicker tone related hyperactivity after 1 s of notched noise presentation (Norena et al., 2003).

According to our model, the increased neural noise to the DCN which is necessary for SR is supposed to originate from the somatosensory system (Krauss et al., 2016, 2018, 2019b). In analogy to the afore mentioned phenomenon of tinnitus modulation by voluntary jaw movements, our model also predicts a modulation of the Zwicker tone perception by somatosensory stimulation. It has indeed been reported that transcutaneous electrical stimulation has an effect on Zwicker tone perception (Ueberfuhr et al., 2017).

## Residual Inhibition

In 1971 Feldmann found that the presentation of acoustic noise leads to a suppression of the tinnitus precept after noise offset (Feldmann, 1971), for approximately one minute (Roberts et al., 2006, Roberts, 2007). This effect was named Residual Inhibition (RI; Vernon, 1977, Henry & Meikle, 2000).

RI should not be mixed up with tinnitus masking, where tinnitus is perceived less intense as it is masked by a noise of similar frequency range (Hazell & Wood, 1981, Terry et al., 1983). In contrast, the presentation of masking noise causes RI *after* the end of noise presentation. As RI is a technique to temporarily modulate the tinnitus percept, it is a potential target for experimental studies on tinnitus mechanisms (Deklerck et al, 2019).

Interestingly, it was reported that RI works best when the masking noise covers the range of the hearing loss of the subjects and is related to the tinnitus pitch (Roberts et al., 2006, 2008). The cause of the suppression of the tinnitus percept during RI has been discussed to be a decreased spontaneous neural activity after masking noise offset (Galazyuk et al., 2017). This is in line with the explanation that there is a neural adaptation along the auditory pathway induced by the noise presentation (Fournier et al., 2018).

These findings emphasize the idea that spontaneous activity of spiking neurons or in other words internally generated neural noise are crucial for processing of acoustic stimuli along the auditory pathway (Galazyuk et al., 2019). This internal noise is suppressed after the presentation of external acoustic noise. To understand the basic neural mechanisms of RI as well as auditory phantom perception, it is crucial to gain a better understanding of how the neural noise contributes to auditory processing.

The idea that the neural system exploits the effect of SR to improve hearing (Krauss et al., 2016, 2018, 2019b) provides a putative explanation for the effect of RI. As described above, tinnitus is potentially induced by the deprivation of neurons along the auditory pathway in tonotopic regions where a cochlea damage occurred. Thus, the auditory system tries to compensate for this deprivation, i.e. hearing loss, by adding internally generated neural noise. This internally generated noise potentially produced by the somatosensory system and fed to the DCN is propagated along the auditory pathway to the cortex, where it is perceived as auditory phantom percept. RI is potentially the consequence of replacing internally generated neural noise by external acoustic noise. In this view, the external noise would replace the internal noise, thereby causing its downregulation and thus suppression of the tinnitus percept as already described in previous publications (Krauss et al., 2016, 2019b).

According to our model, the optimal noise is tuned and controlled on time scales of seconds via a control circuit (Krauss et al., 2016; Figure 2). From this point of view, Zwicker tone and tinnitus are basically the same phenomenon, but on different time scales. Furthermore, the proposed control circuit would work inversely for Zwicker tone and RI. Whereas, the Zwicker tone corresponds to an upregulation of internal neural noise caused by a reduced auditory input (i.e. the notch), RI in contrast corresponds to a downregulation of internal noise, due to increased auditory input (i.e. external acoustic noise). Thus, both phenomena can be considered to be opposite effects that may be explained by exactly the same neural control circuitry proposed by our SR model. To put it in a slogan, the SR model of auditory processing suggests that “RI can be interpreted as an inverse Zwicker tone illusion”.

## Summary and Discussion

In summary, our SR model provides a unified explanation for the induction of acute subjective tinnitus, Zwicker tone, and RI. Especially a look at the time scales, in which Zwicker tone (cf. Norena et al., 2003) can be induced or tinnitus can be reduced by hearing aids or cochlear implants (McNeill et al., 2012, Ito & Sakakihara, 1994, Baguley & Atlas, 2007), indicates that these phenomena cannot be exclusively explained by brain plasticity. The SR model, describing tinnitus as a side effect of the neural system trying to optimize information transmission after hearing loss by exploiting the SR effect, would offer an explanation of how these phantom perceptions can be induced or suppressed so quickly. Thus, the neural system does not need any plasticity as the SR mechanism is optimized by a simple control circuit (Krauss et al, 2016; Figure 2).

We speculate that in subjects, where the Zwicker tone can be induced by short noise presentation the RI effect should vanish more quickly, because the tuning of the optimal noise level works faster in certain subjects and thus the downregulated neural noise during RI is quickly re-increased. On the other hand, the Zwicker tone is induced faster as the neural noise is quickly upregulated when notched noise is presented. Thus, the duration of notched noise needed to induce the Zwicker tone could potentially correlate with the duration of the RI effect. This would be only the case, if both effects were produced by the same SR control circuit in the DCN (Figure 2), which could be a characteristic feature of different individuals. The characteristic parameter of this control circuit is the time needed for controlling the noise amplitude.

This is a testable hypothesis derived from the SR model, which has to be verified or falsified in future studies.

However, it is obvious that the SR model has some limitations, such as that -in contrast to homeostatic plasticity models- it does not predict massive structural and functional changes (cf. Norena, 2011) along the auditory pathway, which is indeed found in several studies (Yang et al., 2011, Li et al., 2015, Singer et al., 2013). These findings are supported by computational models demonstrating the influence of this plasticity (Schaette & Kempter, 2006, Nagashino et al., 2012).

Additionally, our model does not address the question why not all people with hearing loss perceive or even suffer from tinnitus. The influence of stress (Mazurek et al., 2012, 2015) and psychological burden (Landgrebe & Langguth 2011; Langguth et al., 2007, 2011) on tinnitus percepts was shown in several studies. Furthermore, the model does not differentiate between chronic and acute tinnitus.

Despite these limitations, we are convinced that we now have the knowledge to draw a complete picture in the light of preceding studies. Figures 3 and 4 provide an overview of the main models and their explanatory power for tinnitus development and Zwicker tone perception. The different models work on different time scales, as well as in different brain areas, as illustrated in Figure 5.

**Figure 3:**
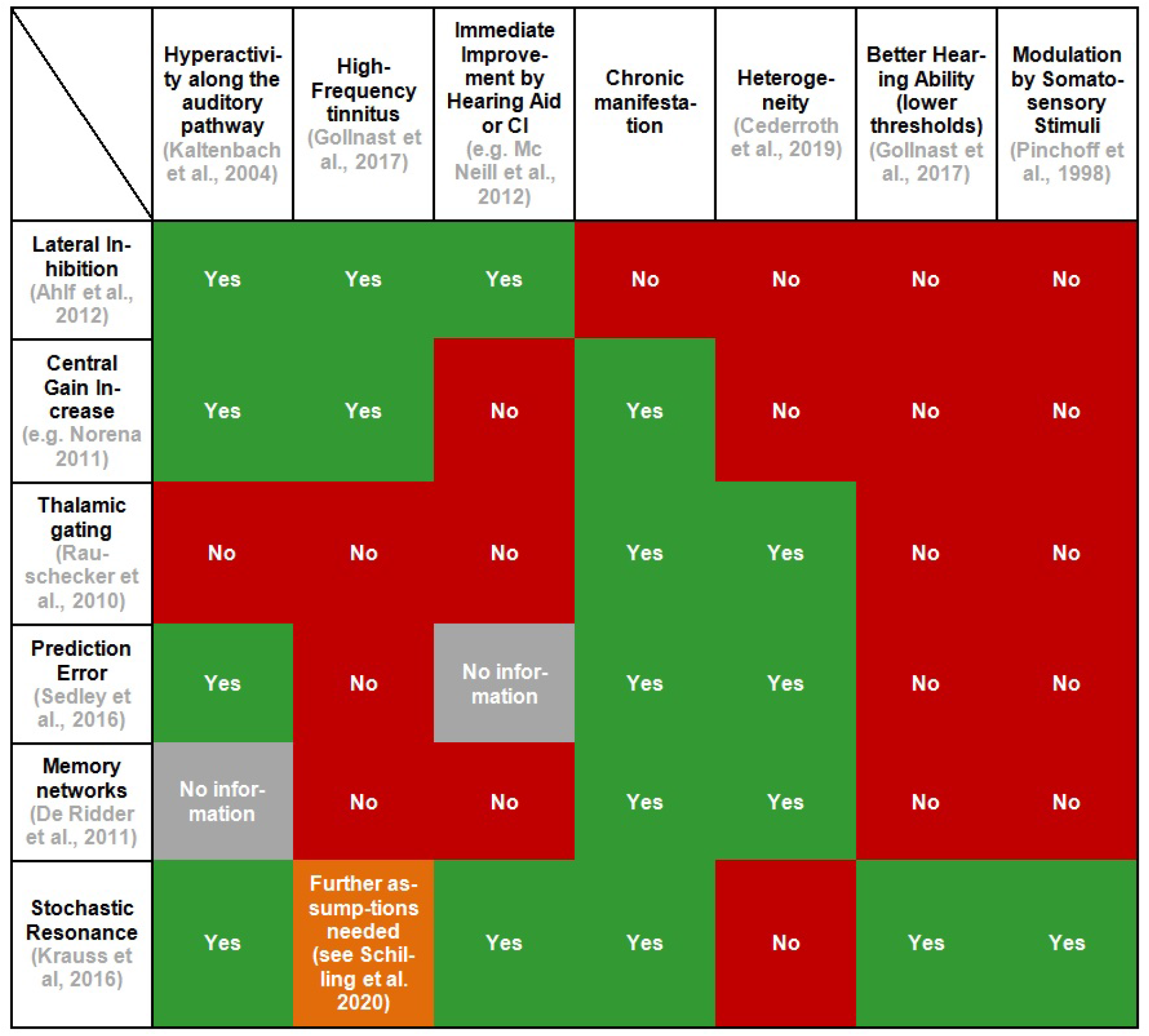
Explanatory power of different models of tinnitus development. The figure summarizes different models of tinnitus development (rows) and how these models fit to certain observations (columns). For each model and effect, one exemplary paper is cited.

**Figure 4:**
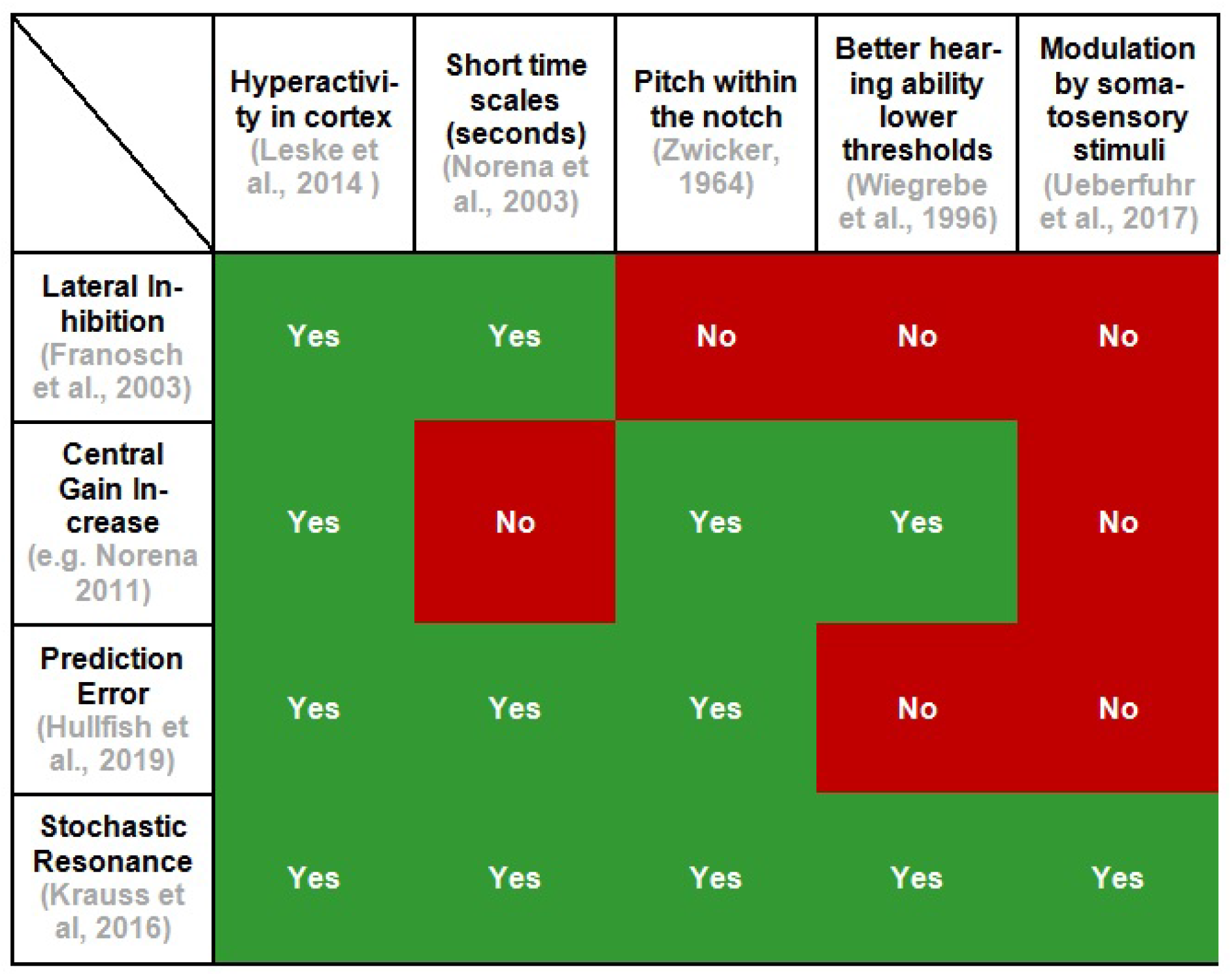
Explanatory power of different models of the Zwicker tone illusion. The figure summarizes different models of the Zwicker tone illusion (rows) and how these models fit to certain observations (columns). For each model and effect, one exemplary paper is cited.

**Figure. 5:**
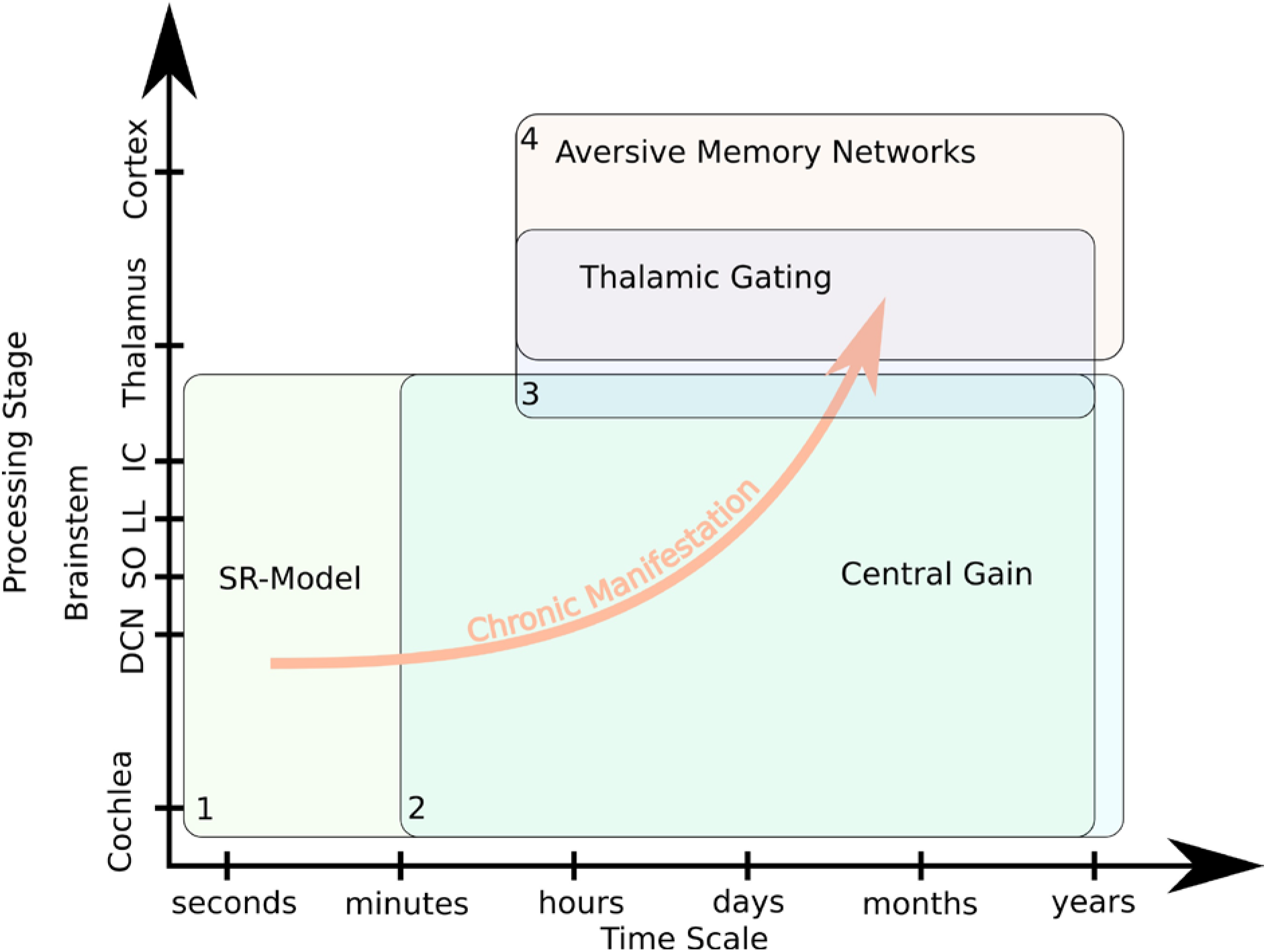
The space of tinnitus models. Models of tinnitus development can be defined at different levels of description and can vary in time scale of the explained observations (horizontal axis) and in proposed anatomical substrate, i.e. processing stage (vertical axis). The SR model fills the “missing gap” in time scales of minutes and seconds.

Our SR model provides an explanation for the induction of tinnitus after e.g. a loud acoustic noise presentation, the induction of Zwicker tone by bandpass noise, or the suppression of the tinnitus percept by acoustic noise presentation (RI). These mechanisms are very fast and occur within seconds, and thus cannot be explained by any of the models based on brain plasticity. However, as described above, some plasticity can be found along the auditory pathway (Yang et al., 2011, Li et al., 2015, Singer et al., 2013). This plasticity could potentially be the first step in chronic manifestation of the tinnitus percept. However, it is still unclear why the gating function of the thalamus does not prevent the neural hyperactivity from being directly transmitted to the cortex as it does for other unwanted permanent stimuli (McCormick & Bal, 1994). This effect could be explained by the model of Rauschecker and coworkers (Rauschecker et al., 2010). There, the auditory input can be cancelled out by the medial geniculate nucleus within the thalamus. This noise cancellation function can be modulated by the limbic system especially the nucleus accumbens, which is indirectly connected to the medial geniculate nucleus. A breakdown of this system impairs the gating function of the medial geniculate nucleus (Rauschecker et al., 2010) and thus brings the neural hyperactivity to consciousness.

De Ridder and coworkers go even one step further and assume a conscious tinnitus percept to be a consequence of different overlapping brain networks including pre-frontal areas as well as brain structures responsible for emotional labeling of certain memories such as the amygdala. Thus, learning effects are involved, which generate a connection of the phantom percept and distress (De Ridder et al., 2011). Unfortunately, this model does not provide mechanistic explanations at a neural network level, but it explains the involvement of different brain structures. Nevertheless, the model could provide an explanation why not every hearing loss causes tinnitus, and why not everyone perceiving tinnitus also suffers from it. Individual memories and neuronal pathways could lead to different effects in different subjects.

The described models draw a complete and consistent image of tinnitus development, chronic manifestation, and heterogeneity, and do not “contradict” each other as described by Sedley and coworkers (Sedley et al., 2016). Furthermore, mechanistic explanations for RI, Zwicker tone, and better hearing thresholds of tinnitus patients compared to patients without tinnitus (Krauss et al., 2016, Gollnast et al., 2017) support the model.

